# Non-invasive prediction of conduction velocities in the human brain from MRI-derived microstructure features at 7 Tesla

**DOI:** 10.1101/2025.10.28.685017

**Authors:** Saina Asadi, Arthur Spencer, Yasser Alemán-Gómez, Maciej Jedynak, Erick J. Canales-Rodriguez, Tommaso Pavan, Emeline Mullier, Louise de Wouters, Isotta Rigoni, Hélène Lajous, Michael Chan, Alexandre Cionca, Dimitri Van De Ville, Serge Vulliémoz, Olivier David, Patric Hagmann, Ileana Jelescu

## Abstract

The conduction velocity of neuronal signals along axons is a key neurophysiological property that can be altered in various disease processes. While cortico-cortical evoked potentials (CCEPs) can be measured in presurgical assessment to provide information about conduction delay between a subset of brain regions, it is currently not possible to efficiently and systematically estimate conduction velocity *in vivo* across the whole brain.

Given the established link between conduction velocity and axon morphology (most notably axon diameter but also myelination), mapping a reliable and quantitative metric linked to axon properties could fill the gap of inferring conduction velocity across the entire human brain. By integrating multiple MRI-derived microstructural measures – including axon radius, axonal water fraction, extra-axonal perpendicular diffusivity, and longitudinal relaxation time – and conduction velocity estimates obtained from a large database of CCEPs, we developed a whole-brain prediction model of conduction velocity. Our multivariate MRI-based model explained 29% of variance in neurophysiological conduction velocity, making it possible to partially predict whole-brain conduction velocity and delay matrices along connections for which no direct measurement is commonly available from epilepsy surgery investigations. This integrative MRI-based approach could provide a non-invasive framework for comprehensively characterising conduction delays *in vivo* across the human brain white matter.

## Introduction

Neuronal delays play a key role in brain dynamics as they encode causal interactions and directional communication pathways between brain areas. These delays arise primarily from the conduction velocity of signals along axons, which ranges widely from 0.5-2 m/s in small, unmyelinated fibres to 50-90 m/s in large, heavily myelinated tracts; with intermediate velocities (3-25 m/s) in cortical and subcortical fibres (Feher, 2012; Hursh, 1939; Purves et al., 2001; Ritchie, 1982; Swadlow et al., 1978; Waxman, 1980). Conduction velocity in the brain is primarily determined by the axon diameter and myelination (Rushton, 1951; Waxman & Bennett, 1972), with myelin promoting rapid saltatory conduction (Hartline & Colman, 2007).

Currently, conduction delays are primarily inferred using invasive electrophysiology in animals, selective invasive measures in humans (during surgery), non-invasive neuroimaging and evoked potentials (Caminiti et al., 2013; Innocenti et al., 2014; Lemaréchal et al., 2022; Trebaul et al., 2018; van Blooijs et al., 2023). However, accurately estimating these delays non-invasively is challenging due to limitations in spatial resolution, difficulties distinguishing axonal conduction delays from synaptic integration times, and the sparse coverage of direct invasive methods.

In a recent study, Lemaréchal et al., (2022) proposed an *in vivo* estimation of the synaptic time constants and axonal conduction delays between brain regions of the human cortex by leveraging a large database of cortico-cortical evoked potentials (CCEPs). The CCEPs are measured invasively using stereo-electroencephalography (SEEG), where intracerebral depth electrodes are employed to stimulate and record neural activity, notably to outline seizure onset zones and epileptogenic networks (Bartolomei et al., 2017; David et al., 2010; Tomlinson et al., 2025; Wendling et al., 2010). The conduction delay between the stimulated and recorded areas is estimated from the measured delay of the first component of the CCEP. If the distance between the two areas is known (e.g., from magnetic resonance imaging (MRI) tractography), then the conduction velocity is inferred from the ratio between distance and delay. While CCEPs provide valuable directional delay connectivity matrices highlighting asymmetric reciprocal connections, their utility at high spatial resolution is limited by sparse sampling and insufficient coverage of the whole brain due to lack of direct connections or an insufficient number of significant responses with accurate fitting. Furthermore, these intracerebral measurements are limited to patients who undergo surgical interventions for epilepsy. To overcome this limitation, MRI-based microstructural measures can offer comprehensive, non-invasive estimations of conduction delays by quantifying key determinants of conduction velocity in the brain white matter (WM), including axon diameter, density and myelination (Drakesmith et al., 2019; Moore et al., 1978).

Diffusion MRI (dMRI) sensitises the MR signal to the random motion of water molecules, which is influenced by the microstructural organisation of brain tissue. Inferring specific cellular features at the sub-voxel level requires combining dMRI acquisition protocols with biophysical models (Alexander et al., 2019; Novikov et al., 2019). Biophysical models are a valuable tool for neuroscience and clinical applications by providing not only sensitivity but also specificity to different physiological and pathological processes (Jelescu et al., 2020). This has been highlighted in several applications including aging (Fan et al., 2019; Toschi et al., 2020), dementia (Dong et al., 2020; Vogt et al., 2020), multiple sclerosis (Bagnato et al., 2019; Liao et al., 2024), ischemia (Liao et al., 2024) and schizophrenia (Pavan et al., 2025).

The standard-model imaging (SMI) framework (Novikov et al., 2018) models the diffusion signal in WM as stemming from the weighted contribution of: 1) an intra-axonal space where axons are modelled as impermeable zero-radius cylinders (known as sticks) arranged in locally coherent fascicles; 2) an extra-axonal space of each fascicle which is modelled with an axially symmetric diffusion tensor; and 3) the cerebrospinal fluid (CSF), an optional compartment modelled with an isotropic tensor. In the WM, the myelin sheath surrounding axons forms compartment “impermeability” over the diffusion timescale (20 to 100 ms), making inter-compartment exchange negligible. SMI parameters are thus: intra-axonal diffusivity (D_a_), extra-axonal parallel and perpendicular diffusivity (*D*_e,∥_, *D*_e,┴_), axonal water fraction (*f*), fibre alignment (*p*_2_) and optionally the CSF water fraction (*f*_iso_; Coelho et al., 2022; Novikov et al., 2018).

Each SMI parameter is specific to a certain feature of the microstructure. For example, *f* measures the relative contribution of intra and extra axonal water, making *f* a good marker of axonal density, and to a lesser extent of myelination (Jelescu et al., 2016). *D*_*a*_ evaluates axonal integrity and beading (Budde & Frank, 2010; Lee et al., 2020). *D*_e,┴_ is highly sensitive and quite specific to myelination which makes the extra-axonal space more tortuous (Jelescu et al., 2016). Finally, *D*_e,∥_can detect broader extra-axonal changes related to extra-axonal crowding, e.g. neuroinflammatory processes like astrogliosis or microglial activation (Jelescu & Fieremans, 2023; Xie et al., 2010). Based on the relationship between conduction velocity and the underlying microstructure features, the SMI parameters that are expected to be most tightly associated with conduction velocity are thus *f*, *D*_e,┴_ and *p*_2_.

In addition to these microstructure metrics, axon diameter estimation is particularly compelling due to its direct correlation with conduction velocity, alongside myelination (Hursh, 1939; Rushton, 1951; Waxman, 1980; Waxman & Bennett, 1972). In a sensitivity analysis based on simulations of axonal physiology, axon radius and g-ratio (i.e., inner to outer axon radius ratio) emerged as the primary determinants explaining variability in conduction velocity across a wide range of parameters (Drakesmith et al., 2019). However, estimating metrics associated with axon radius distributions from MRI is challenging. For acquisition parameters achievable on clinical scanners, the axons can effectively be modelled as sticks, i.e., cylinders with zero radius, as in SMI (Veraart et al., 2018). Advanced hardware with strong field gradients, such as Connectome scanners (Veraart et al., 2019), can improve sensitivity to axon diameter (Duval et al., 2015; Veraart et al., 2020). Nevertheless, the dMRI sensitivity to axon diameters is intrinsically limited by the dependence of the diffusion-weighted signal in cylinders to their radius distributions, whereby the effective estimated radius varies as 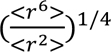. In other words, the estimated axon radius from dMRI is heavily skewed towards the tail of the distribution (Veraart et al., 2020), as illustrated in initial – ten-fold overestimated – dMRI axon diameter maps (Alexander et al., 2010; Assaf et al., 2008; Dyrby et al., 2013). Histological studies have reported axon diameters in human WM to range from 0.5 to 2 µm (Aboitiz et al., 1992a; Liewald et al., 2014), while only 1% of all axons have diameters larger than 3 µm (Caminiti et al., 2009).

A novel diffusion-relaxation MRI method was recently proposed, which relies on T_2_ surface relaxation in the intra-axonal space instead (Barakovic et al., 2023). This ap roach addresses the challenge of measuring small axon radii in the brain using MRI, because here the effective estimated radius 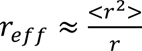 is closer to the mean of the axon radius distribution <r> than those provided by pure dMRI models which are heavily weighted by the tail of the distribution. This approach relies on diffusion-weighting only to preferentially filter out the extra-axonal signal by applying sufficiently high *b*-values (i.e., b > 5-6 ms/µm^2^). This promising method was tested using postmortem histological measurements (Barakovic et al., 2023) and validated in axon-mimicking microfibers phantoms (Canales-Rodríguez et al., 2024). A significant advantage of this method is that, unlike pure dMRI models, it could be implemented on human clinical scanners, as it does not require the use of ultra-strong diffusion gradients that are only available on preclinical scanners or human “Connectome” scanners.

Finally, myelin content influences conduction velocity in the WM, which can also be captured using complementary MR contrasts (Piredda, Hilbert, Thiran, et al., 2021). For example, the longitudinal relaxation rate (R_1_ = 1/T_1_) in the WM has been associated with myelin content, as water interacting closely with myelin exhibits shorter T_1_ relaxation times compared to free water (Lutti et al., 2014; Mezer et al., 2013; Stüber et al., 2014). However, 1/T_1_ has also been reported to correlate with axon radius. Since T_1_ reflects relaxation across both intra and extra-axonal compartments, the association with axon diameter may in fact be stronger (a weighted measure of inner and outer radii), though influenced by additional confounding factors (Harkins et al., 2016; Hofer et al., 2015).

Testing microstructural features against conduction velocity is challenging, as there is no non-invasive, whole-brain ground truth for conduction velocity in humans; direct validation requires invasive stimulation and extensive sampling. The Functional Tractography (F-TRACT) project (Avalos-Alais et al., 2025; Jedynak et al., 2023; Trebaul et al., 2018) addresses this by integrating large-scale SEEG single-pulse electrical stimulation data from hundreds of patients to construct a probabilistic atlas of cortico-cortical connectivity and CCEP-derived delays. Correlating the MRI-derived microstructural measures with the electro-neurophysiological atlas of conduction delay (F-TRACT) can quantify their association and enable whole-brain mapping of neuronal conduction delays. Based on these associations, we develop an MRI-based model, validated against CCEP data, to non-invasively predict conduction velocity and delays throughout the human WM *in vivo*, including in connectivity edges for which F-TRACT data is not available.

## Materials and Methods

### Participants and demographics

The study was approved by the ethics committee of the Canton of Geneva. All participants signed written informed consent. Healthy participants (N = 20; females = 10; median age = 25 years, interquartile range (IQR) = 30-23 years, range = 21-65 years) were scanned on a clinical 7T MR system equipped with 120 mT/m gradients (MAGNETOM Terra.X, Siemens Healthineers, Erlangen, Germany) using an 8 channel transmit and 32 channel receive head coil.

### Image acquisition

**Structural MRI:** T_1_-weighted 3D anatomical images were acquired using an MP2RAGE sequence. Acquisition parameters were as follows: repetition time (TR) = 6000 ms; echo time (TE) = 2 ms; TI_1_ = 800 ms (4° flip angle, inv1) and TI_2_ = 2700 ms (5° flip angle, inv2); spatial resolution = 0.64×0.60×0.60 mm³; field of view = 164×236×250 mm^3^; matrix size = 256×392×416; acquisition time (TA) = 7 minutes. The quantitative T_1_ map was computed from the images at the two inversion times.

**Diffusion MRI:** Diffusion-weighted (DW) images were acquired using a 2D multi-slice PGSE-EPI sequence with parameters as follows:

***Multi-shell DWI:*** b [ms/µm²] (directions) = 0 (4), 1 (20), 2 (40), 3 (60); δ = 10.6 ms; Δ = 39.6 ms; TE = 78 ms; TR = 5000 ms; 1.5-mm isotropic resolution; field of view = 230×230×120 mm^3^; matrix size = 152×152; 80 slices; acceleration = GRAPPA 3 x multiband 2; TA = 11 minutes.

***Multi-TE DWI:*** b [ms/µm^2^] (directions)=0 (4), 5 (64), TR = 5000 ms, 2-mm isotropic resolution; field of view = 230×230×120 mm^3^; matrix size = 114×114; 60 slices; acceleration factor = 6 (9 subjects: GRAPPA 3 x multiband 2; 11 subjects: GRAPPA 2 x multiband 3). The sequence was acquired for four different echo times: TE = 80, 100, 120, 135 ms, TA = 25 minutes.

A b = 0 image with reversed phase-encoding direction was acquired in both multi-shell and multi-TE acquisition for geometric distortion correction.

### Image pre-processing

DW images were pre-processed using the following pipeline: noise level estimation and removal using magnitude and phase data with NOise Reduction with DIstribution Corrected (NORDIC) principle component analysis method (Moeller et al., 2021) implemented in MATLAB (*version R2021b* or newer); Gibbs ringing correction using “mrdegibbs” in MRtrix3 (*version 3.0.4*; Kellner et al., 2016; Tournier et al., 2019); susceptibility and eddy current distortion correction using “top-up” and “eddy” (J. L. R. Andersson et al., 2003; J. L. R. Andersson & Sotiropoulos, 2016) tools in FSL (*version 6.0.7.11*). For the multi-TE data, an additional motion correction step was applied per TE to further remove motion across all the volumes. Specifically, we first estimated the diffusion tensor model from the eddy corrected data. Each DW volume was then registered to its corresponding model-predicted volume using ANTs rigid body registration (*version 2.4.4*; Avants et al., 2011). From the motion-corrected data, we re-estimated the diffusion tensor model and corresponding signal, followed by a second registration step to further reduce residual misalignment (Figure 1).

**Figure 1.**
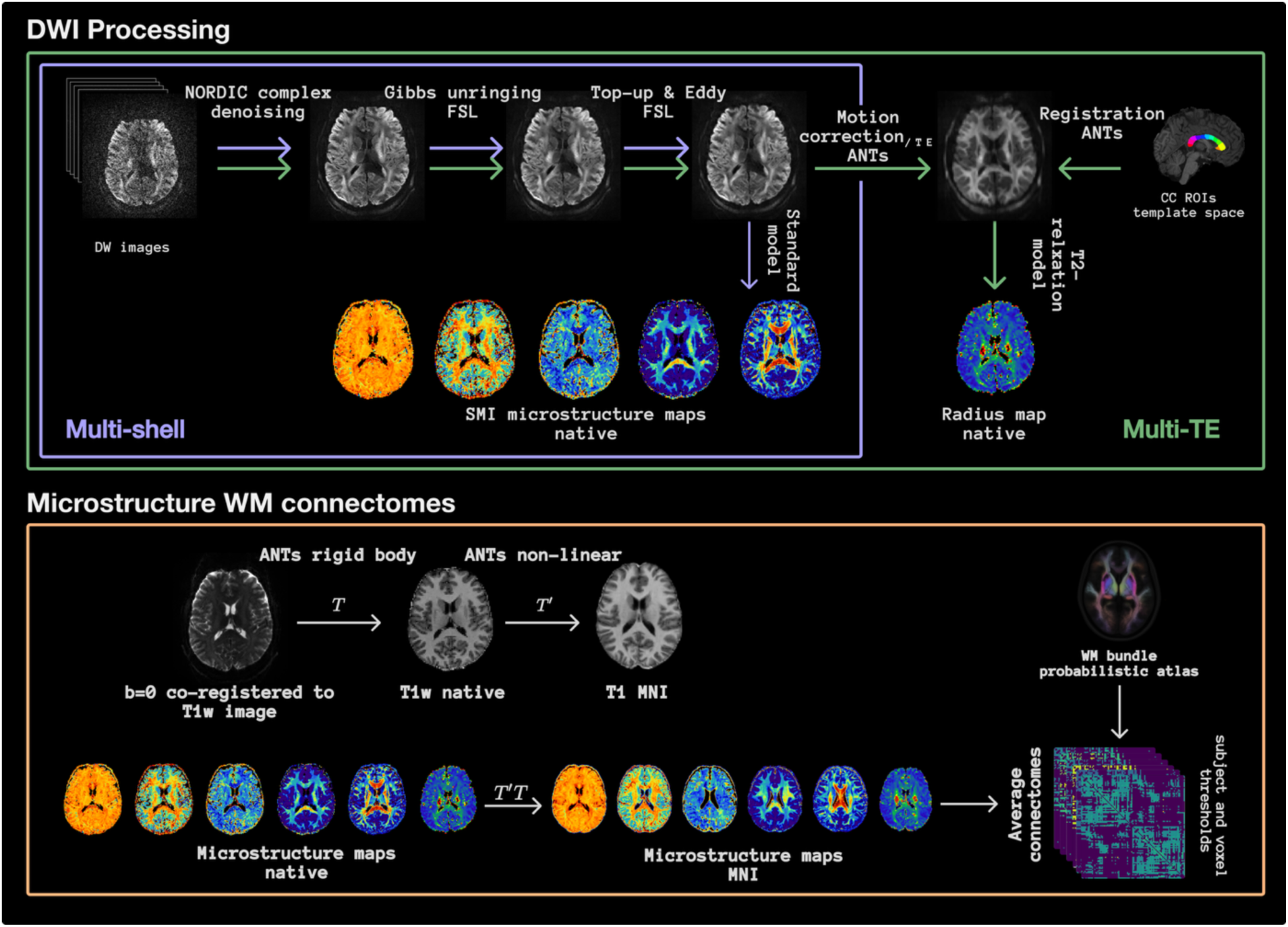
Processing of diffusion-weighted data, from diffusion images to microstructure connectomes. The pre-processing steps of the multi-shell and multi-TE data are illustrated with the purple and green representations, respectively. All the microstructure parameters are estimated in subject space, then registered to standard MNI space for the connectivity estimation within the scale 1 MultiConn atlas of 95 ROIs (Alemán-Gómez et al., 2022). DWI = Diffusion Weighted Image, CC = Corpus Callosum, SMI = Standard-Model Imaging, WM = White Matter.

### Standard Model Imaging (SMI) parameters

WM microstructure parameters were estimated from the multi-shell DW data using the SMI framework (Coelho et al., 2022; Novikov et al., 2018). Quantitative maps of intra-axonal diffusivity (*D_a_*), parallel and perpendicular extra-axonal diffusivities (*D*_e,∥_, *D*_e,┴_), axonal water fraction (*f*), and fibre alignment (*p*_2_) were computed for all subjects. Note that in typical multi-shell protocols with single TE, introducing a third compartment for the CSF will further increase the difficulty of estimating other SMI parameters (Coelho et al., 2024; Liao et al., 2024), therefore we only considered a two-compartment model, without the CSF. The SMI microstructure maps were registered to standard MNI space for further connectivity analyses; first, the b = 0 image was co-registered to the native T_1_-weighted image, and the T_1_-weighted image was registered to MNI space using ANTs non-linear registration (*version 2.4.4–*SyNCC method; Avants et al., 2011). The combined transformation was then applied to the microstructure parametric maps. Finally, the group averaged parametric maps were computed across subjects.

### Axon radius

Axon radii were estimated from the DW multi-TE data by applying a diffusion-relaxation model, as outlined by Barakovic et al., (2023) and Canales-Rodríguez et al., (2024). This model was fitted to the directional average (i.e., spherical mean) of the dMRI signal, as a function of TE. Eventually, only the first three TEs (i.e. 80, 100, 120 ms) were used, due to low signal-to-noise ratio (SNR) of the images acquired with the highest TE.

Data from two histological samples were used to calibrate and test the axon radius estimation, respectively (Barakovic et al., 2023). The first sample, containing axon radius measurements from four regions of interest (ROIs) within the corpus callosum (CC), served to calibrate the T_2_-based model parameters (Figure 2A). Then, the calibrated model was applied across WM to estimate voxel-wise axon radii *in vivo.* The second sample provided histological radius measurements from 11 additional ROIs in the CC and was used as a test set to evaluate the estimation performance of the calibrated model (as a quality control step) in these ROIs. For conducting this analysis, the segmented CC regions were non-linearly transformed from the image space where the histological ROIs were originally defined to each subject’s native space using the ANTs registration (*version 2.4.4–*SyNCC method; Avants et al., 2011). Subsequently, a linear regression model was used to compare the estimated effective axon radii and the histological values (Figure 2B).

**Figure 2.**
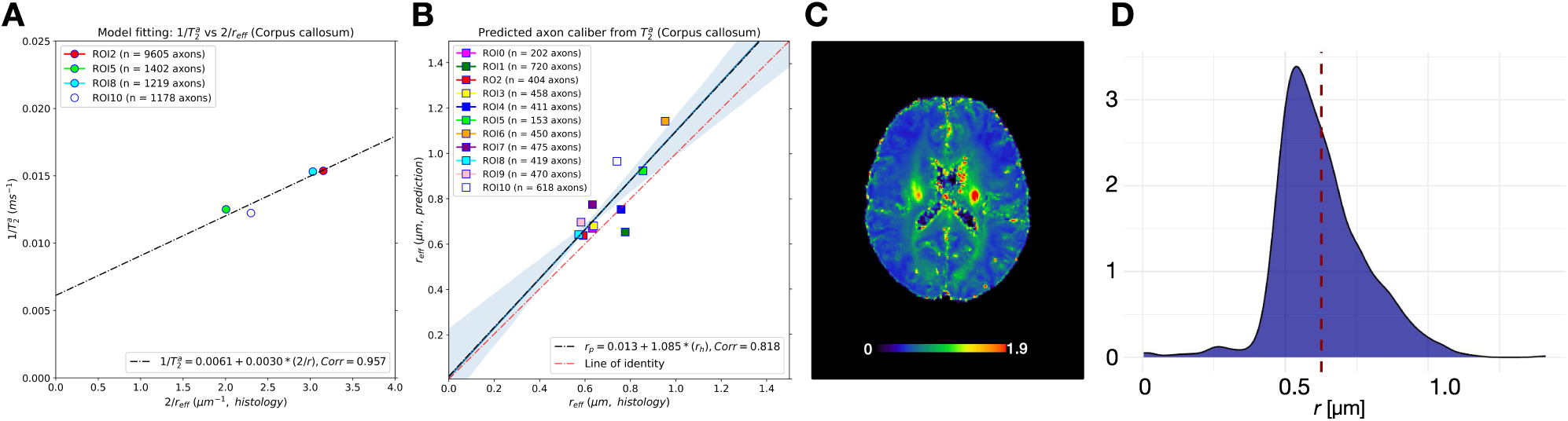
Axon radius estimation. From left to right: **A:** Deriving regression coefficients (calibration step): example linear fitting of the intra-axonal transverse relaxation rate estimated from the *in vivo* multi-TE DW data in one subject to the inverse of the inner axon radius measured from a histological brain sample in four ROIs of the CC. **B:** Test set (quality control step): linear fitting of the predicted radius from the intra-axonal T_2_ relaxation time using calibrated regression parameters from (A) to the effective histological radius estimated in 11 other ROIs of the CC. **C-D:** Average axon radius map and probability density plot of WM axon radius distribution across subjects.

Finally, the axon radius maps were registered to standard MNI space using the same registration procedure described for the SMI parametric maps. A group-level axon radius map was derived by computing the voxel-wise median across subjects to minimize the influence of extreme values from individual subjects. For a detailed acquisition protocol and pre-processing pipeline of the diffusion data, see Figure 1.

### Quantitative T_1_ relaxation time as a myelin proxy

Individual quantitative T_1_ (qT_1_) maps were used as a proxy for myelination (Callaghan et al., 2015; Koenig et al., 1990; Lutti et al., 2014; Piredda, Hilbert, Thiran, et al., 2021; van der Weijden et al., 2021) and were registered to standard MNI space using non-linear registration in ANTs (*version 2.4.4–*SyNCC method; Avants et al., 2011). The group averaged qT_1_ map was then computed across subjects.

### Microstructure Connectomes

We used the probabilistic multi-scale WM bundle atlas (MultiConn atlas) by Alemán-Gómez et al., (2022) which is derived from a cohort of 66 healthy subjects of the Human Connectome Project (HCP; Van Essen et al., 2013). We selected the parcellation comprising 95 grey matter regions. The spatial probabilistic anatomical map of each bundle was generated by binarizing and averaging the tract density images across subjects. We set two consistency thresholds for the atlas: 1) a voxel-wise probability threshold of 0.2 to exclude low-probability voxels; and 2) an inter-subject consistency threshold of 70%, indicating the minimum number of subjects with a non-zero fibre count between a given pair of brain regions. As a result, group level microstructure connectomes were quantified as the mean scalar value along the bundle mask connecting each pair of the grey matter regions. Additionally, a length matrix was obtained, where each entry represents the mean geodesic length of streamlines connecting pairs of regions in the atlas.

### SEEG data and axonal conduction delays estimation

The exact procedure of axonal delays estimation is described in the earlier work by Lemaréchal et al., (2022). This procedure was conducted on the F-TRACT dataset (Trebaul et al., 2018; https://f-tract.eu) that aggregated SEEG recordings from 12 different centres from patients with drug-resistant epilepsy. The signals were recorded at rest, in the awake state. The participants gave consent to undergo this clinical procedure and to share their data for research purposes under ethical guidelines of the Internation Review Board at INSERM (protocol number: INSERM IRB C14-18), in agreement with the declaration of Helsinki. Details on axonal conduction delay estimation can be found in the Supplementary Methods.

The obtained axonal delays were grouped according to the stimulated and the recorded parcels and the mean values were computed for groups with at least 10 values. The parcels were delineated according to Lausanne2008-33 atlas (Hagmann et al., 2008) consisting of 83 ROIs corresponding to the 95-ROI (scale-1) MultiConn WM atlas, with a combined thalamus region instead of distinct thalamic nuclei. Conduction velocity for each bundle was derived by dividing tract length by the corresponding mean axonal delay. Since the microstructure connectomes do not encode directional information, conduction delay and velocity matrices were symmetrised by averaging corresponding entries from the upper and lower triangles. Finally, microstructural properties across the 12 thalamic nuclei were averaged to match the 83 ROIs in the F-TRACT delay matrix for subsequent analyses.

### Statistical and model analyses

The analysis was restricted to cortico-cortical connections, to ensure accurate estimation of conduction delays, thereby excluding all subcortical regions resulting in 68 ROIs. In each bundle, we calculated Spearman’s rank correlation coefficients of WM microstructure parameters (most biologically relevant SMI parameters, i.e., *D*_e,┴_, *f* and *p*_2_) and *r*, qT_1_, conduction velocity, bundle length and conduction delay, to test for monotonic associations. Significant results were corrected for multiple comparisons (FDR *α* = .05).

To determine the optimal set of predictors for conduction velocity, we initially specified a full regression model including all predictors of interest, with log-transformed conduction velocity as the dependent variable:

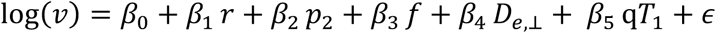

Predictors were then iteratively removed based on statistical significance and model fit criteria until we identified the simplest model that provided optimal explanatory power. A logarithmic transformation was applied to linearise multiplicative relationships between predictors and conduction velocity and to stabilise residual variance, thereby meeting the assumptions of linear regression. The data then were split into 80% training and 20% testing sets, and bootstrap resampling (n = 1,000) was used to estimate model uncertainty and derive error metrics (root-mean-squared error (RMSE), mean absolute error (MAE) with 95% confidence intervals and median absolute percentage error (APE)). The final model was then applied to predict conduction velocity. Delays were predicted from conduction velocities using a bootstrap-based estimation. For each bootstrap iteration, the predicted delay was computed using:

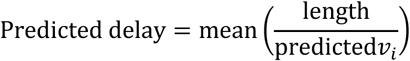

Where *v_i_* is the predicted velocity for bootstrap *i*. The predicted delay was then obtained by averaging the delay estimates across 1,000 bootstrap iterations and the error metrics were computed. Predicted conduction velocity and delay matrices were generated and model uncertainty was evaluated by examining the distribution of uncertainty values.

## Results

### Relationship between the different microstructure metrics

Axon radius estimation was reliable in 16 of the 20 participants; in 4 subjects, motion and artifacts for the multi-TE acquisition prevented an adequate fit during the calibration step (see Methods), and these were excluded from the population average of *r* (Figure 3B). For the SMI parameters and qT_1_, all 20 participants were retained (Figure 3A,C).

**Figure 3.**
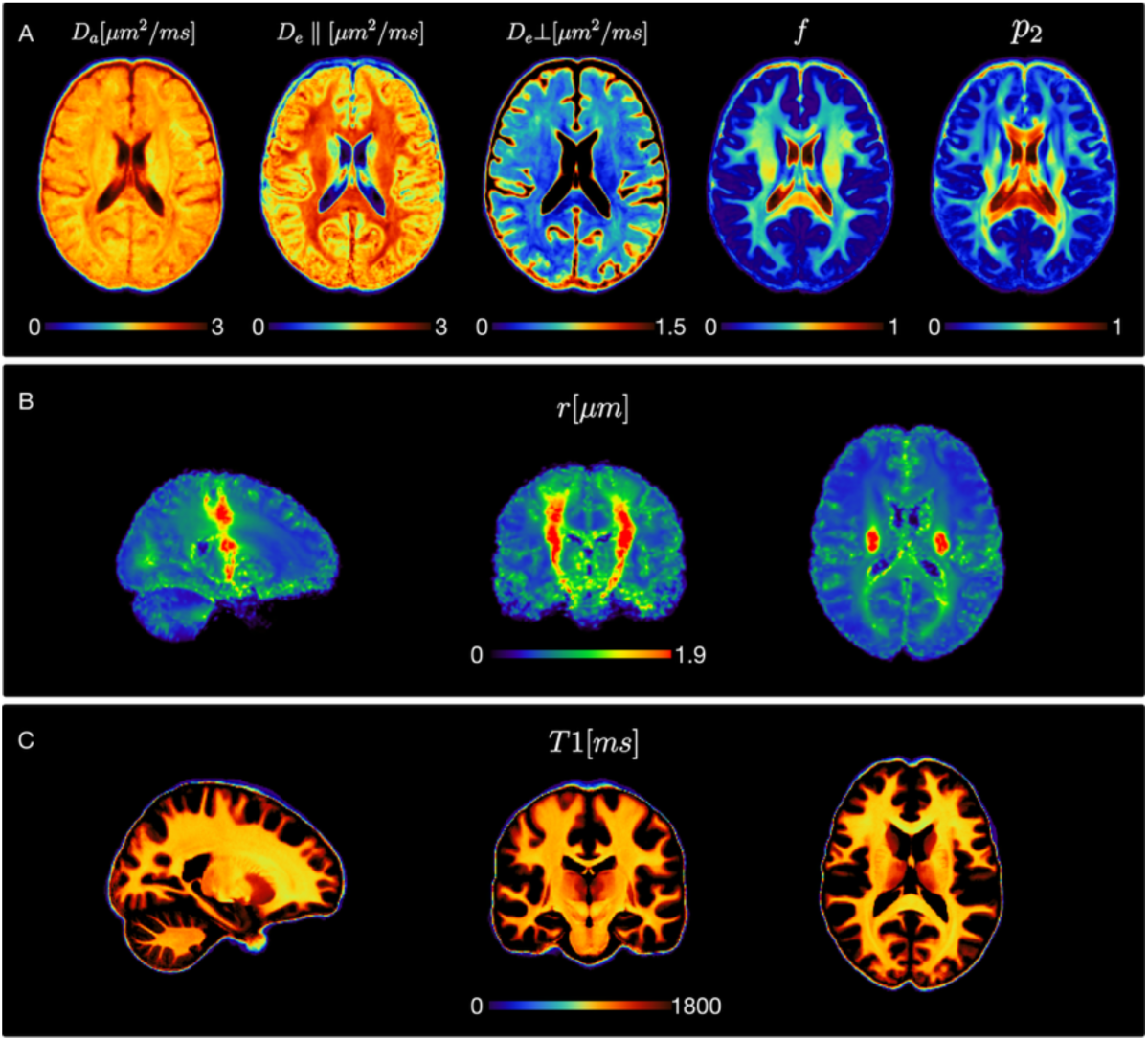
Group averaged parametric maps and atlas-based connectomes. **A:** Columns show different standard-model imaging (SMI) parameter maps for 20 healthy participants. From left to right: intra-axonal diffusivity (D_a_), parallel and perpendicular extra-axonal diffusivities (*D*_e,∥_, *D*_e,┴_), axon water fraction (*f*), and fibre alignment (*p*_)_). **B:** Averaged radius map for 16 healthy participants highlighting the corticospinal tract with highest radius estimates (*r* [Mean (SD)] = 1.38(0.40) µm). **C:** Averaged quantitative T_1_ map for 20 healthy participants.

After deriving the group-average connectomes for the MR-based microstructural metrics, fibre length, and electrophysiological conduction properties (Figure 4), we first assessed the monotonic relationship between different microstructure parameters obtained from DWI, specifically *D*_e,┴_, *f* and *p*_2_, together with *r* and qT_1_. These correlations were performed to validate the consistency of parameter estimates derived from different MR acquisition schemes with established biological relationships.

**Figure 4.**
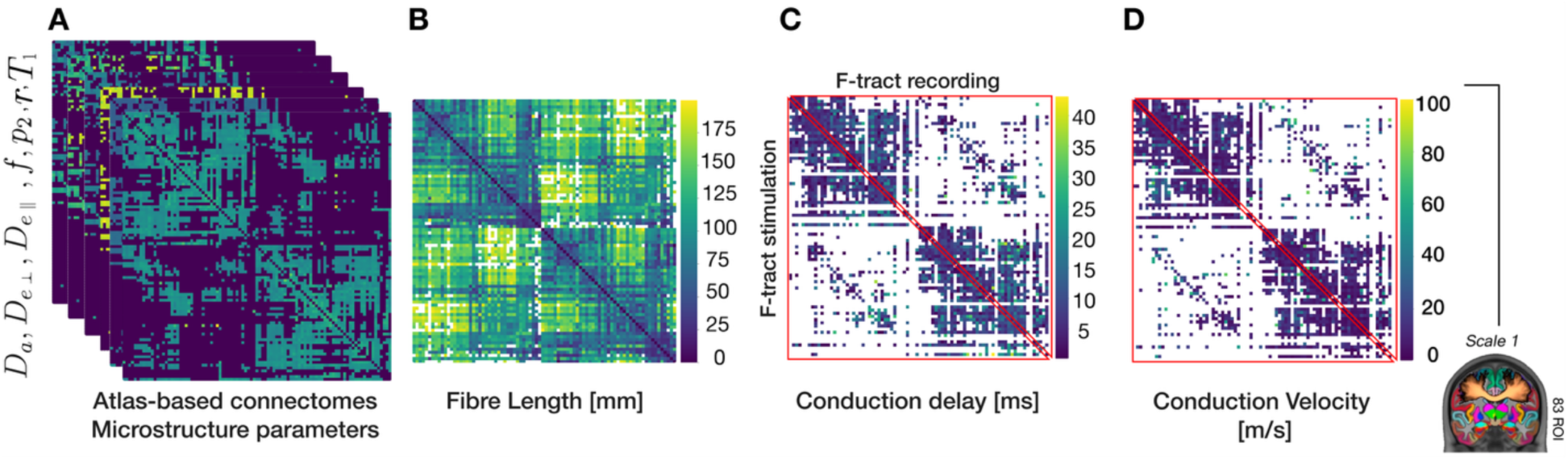
From left to right: **A:** Group averaged microstructure connectomes within the scale 1 WM bundle atlas (excluding the sub-thalamic regions; Alemán-Gómez et al., 2022). **B:** Bundle length matrix derived from the connectome atlas with each entry representing the mean length of the streamlines connecting that pair of regions. **C:** Delay matrix derived from stimulation and recording data from the F-TRACT database (Lemaréchal et al., 2022) corresponding to the ROIs based on the WM bundle atlas. **D:** Conduction velocity matrix computed by dividing the fibre length by the conduction delay matrix; both delay and velocity matrices were then symmetrised by averaging the upper and lower edge values.

The correlation analysis among SMI model parameters indicated moderate to low association, suggesting that the SMI parameter count is adequate to represent the signal within this range without redundancy. In other words, each metric appears to capture distinct information, reinforcing the model’s ability to differentiate between microstructure features. Furthermore, we observed strong positive correlations between *r* and both *f* and *p*_2_ [r = 0.64, *p* < .001; r = 0.58, *p* < .001]. The associations between qT_1_ and diffusion-based microstructure parameters were also prominent, characterized by a positive correlation with *D*_e,┴_ and a negative correlation with *f*, *p*_2_, and *r* [r = 0.51, *p* < .001; r = −0.72, *p* < .001; r = −0.44, *p* < .001; r = −0.21, *p* <.001; Figure 5).

**Figure 5.**
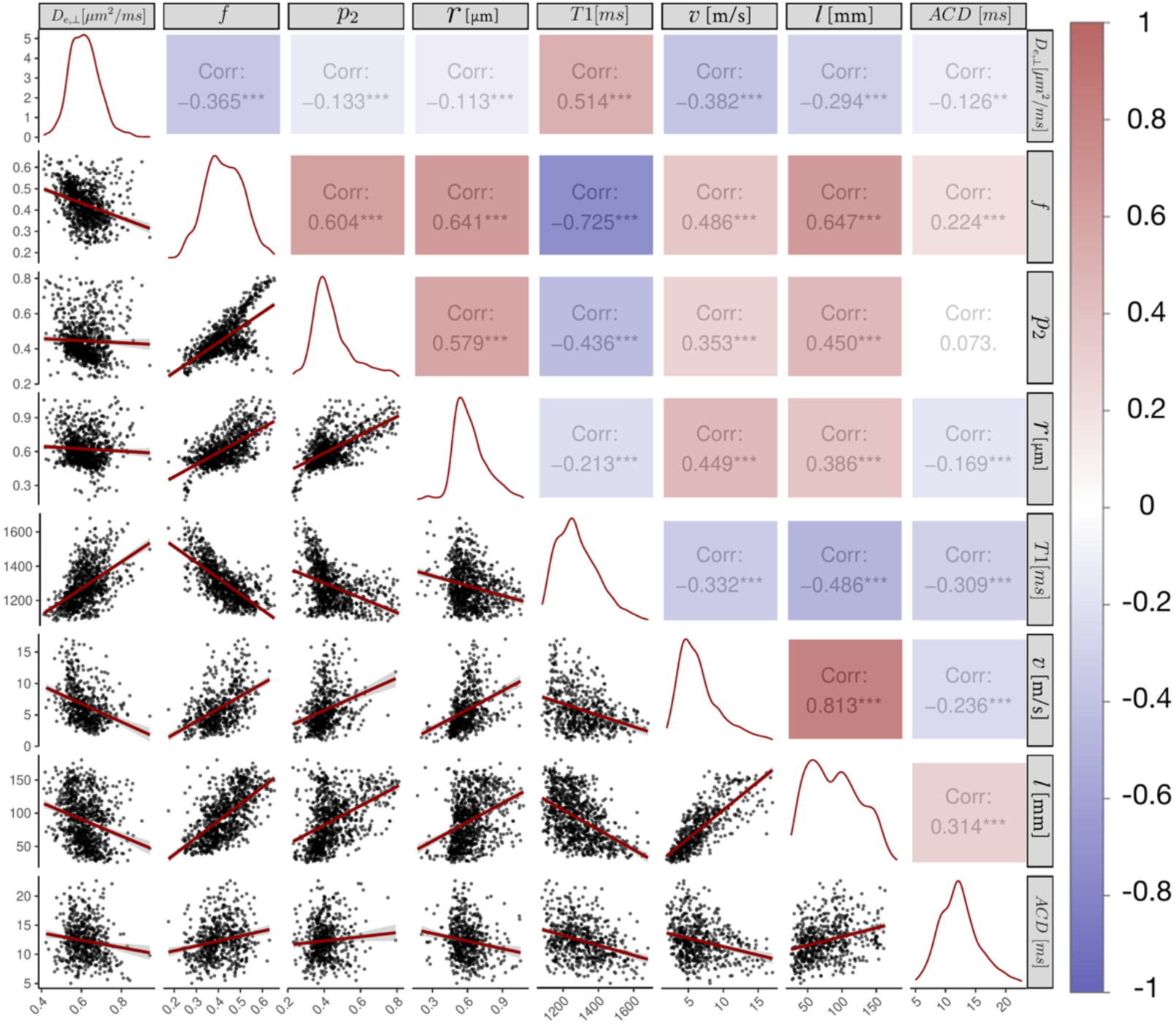
Correlation matrix between dMRI microstructure parameters (i.e. perpendicular extra-axonal diffusivity (*D_e,┴_*), axon water fraction (*f*), fibre alignment (*p*_2_) and axon radius (*r*)) with conduction velocity (*v*), axonal conduction delay (*ACD*) and length (*l*). The diagonal density plots show the distribution of each variable, while the off-diagonal scatter plots illustrate the correlation trends between variables. The upper triangle displays Spearman’s correlation coefficients (displayed *p*-values are FDR corrected; significance at FDR < 0.05), and the lower triangle includes scatter plots with regression lines, providing an overview of the relationships between the variables.

### Microstructure correlates of F-TRACT conduction velocity

The correlation between the microstructure metrics and conduction velocity showed significant positive associations with *f*, *p*_2_ and *r*, respectively [r = 0.49, *p* <.001; r = 0.35, *p* < .001; r = 0.45, *p* < .001; Figure 5], as well as a negative association with *D*_e,┴_ and qT_1_ [r = −0.38, *p* <.001; r = −0.33, *p* < .001; Figure 5]. Our results highlight *f* and *r* as the main microstructure metrics associated with conduction velocity. Note that there was also a strong correlation between conduction velocity and fibre length [r = 0.81, *p* <.001; Figure 5].

### Conduction velocity and delay predictions in the white matter

A multiple linear regression was conducted to predict the conduction velocity from *r*, *f*, qT_1_, and *D*_e,┴_. The final model, reported in Table 1, excluded *p*_2_. This parameter was removed during the model selection as it was not significant after accounting for other microstructural predictors, notably given the collinearity with *r* and *f*.

**Table 1.** Linear model summary. Multiple regression analysis assessing the influence of predictors *r*, *f*, qT_1_, and *D*_e,┴_on the conduction velocity (intercept included). For each term, the unstandardised regression coefficient (β), standardised regression coefficient after the fit (post-hoc standardisation; std. β), standard error (SE), t-statistic, *p*-value, and corresponding 95% confidence interval (CI) are reported.

| Predictor | $\beta$ | $Std. \beta$ | $SE$ | $t$ | $p$-value | 95% CI |
| --- | --- | --- | --- | --- | --- | --- |
| Intercept | 0.13 | NA | 0.42 | 0.30 | 0.764 | [-0.70, 0.95] |
| $r$ | 0.59 | 0.16 | 0.17 | 3.57 | <0.001 | [0.27, 0.91] |
| $f$ | 2.70 | 0.47 | 0.42 | 6.36 | < 0.001 | [1.86, 3.53] |
| $D_{e,\perp}$ | -1.009 | -0.16 | 0.26 | -3.84 | < 0.001 | [-1.52, -0.49] |
| $qT_1$ | 0.0006 | 0.18 | 0.0002 | 2.73 | 0.006 | [0.000, 0.001] |

The overall model was significant, F (4,592) = 62.32, p < .001, and accounted for 29% of the variance in log(*v*). Specifically, *f, r* and qT_1_ showed significant positive associations with log(*v*), whereas *D*_e,┴_ exhibited a significant negative association.

This model was then applied to predict conduction velocity and delay for all connections, including those missing from the F-TRACT connectomes (Figure 6A-B). Predicted velocities and delays closely overlapped with the empirical F-TRACT distributions, both showing positively skewed shapes. Existing F-TRACT connections had moderate velocities (median = 12 m/s) and delays (median = 12 ms), with a longer tail for both measurements. Predictions for missing connections followed the same pattern, indicating physiologically plausible values (Figure 6C).

**Figure 6.**
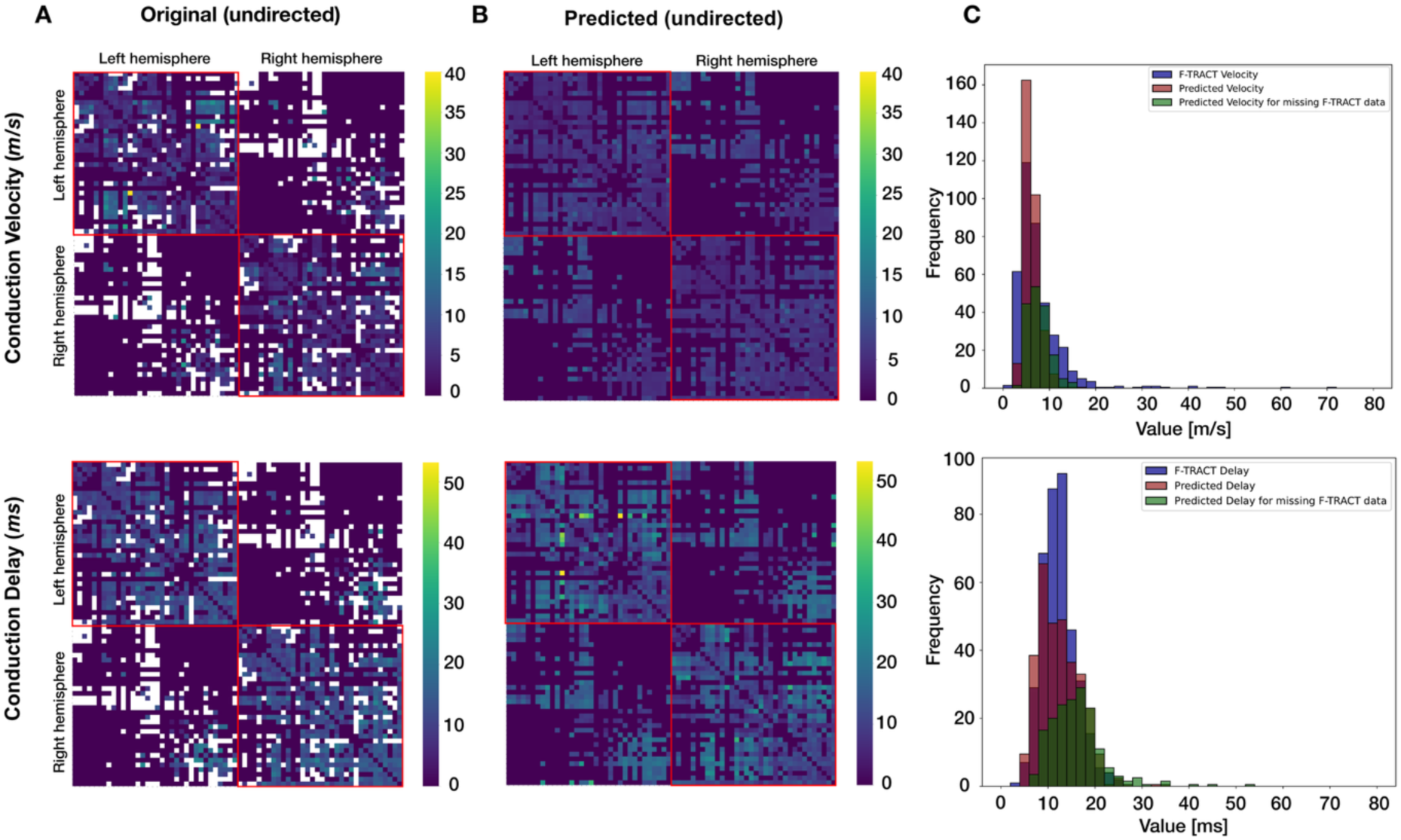
Original and predicted conduction velocity (top) and delay (bottom) matrices and distributions. **A:** Original symmetrised conduction velocity and delay matrices of 68 ROIs (cortico-cortical connections), with missing SEEG cortico-cortical delay recording edges. **B:** Predicted conduction velocity and delay matrices for all connections. The missing connections in the original velocity and delay matrices (white edges) were filled using our model predictions. Entries equal to zero (dark blue) reflect the voxel threshold applied to the MRI parameters’ connectomes (see Methods), which were subsequently matched to the velocity and delay matrices to ensure consistent edge definitions. **C:** Distributions of conduction velocity and delay values across original F-TRACT measurements, MRI-derived predictions, and connections without F-TRACT data.

### Model uncertainty estimates

Model performance was evaluated using bootstrap resampling (n = 1,000) on connections with existing velocity and delay values. Predictions for conduction velocity yielded a RMSE of 2.60 ± 0.13 m/s (95% CI [2.33, 2.85] m/s). In addition, MAE was 1.91 m/s (95% CI [1.75, 2.08] m/s), and the APE was 25%. These results indicate that the model’s predictions are relatively robust and exhibit moderate error variability across bootstrap iterations (Figure 7A).

**Figure 7.**
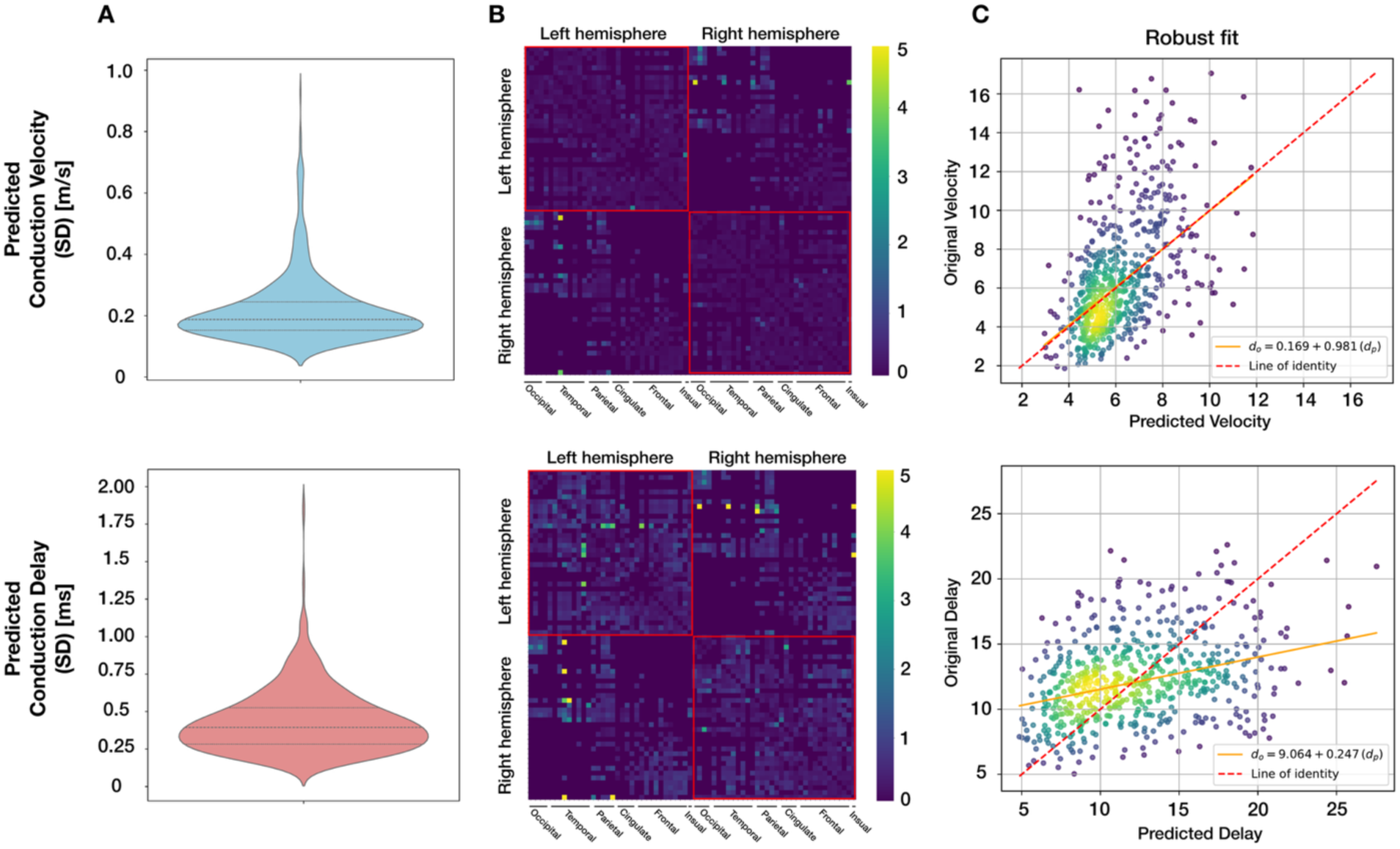
Model uncertainty estimation for conduction velocity (top) and delay (bottom). **A:** Violin plots show the distribution of the model uncertainty for conduction velocity and delay, quantified as the standard deviation of predictions obtained from 1,000 bootstrap iterations for the existing edges. **B:** Heatmap of the 68 ROIs matrix displays the spatial distribution of uncertainty for each connection’s predicted conduction velocity and delay. In both matrices, the uncertainty is computed as the empirical standard deviation across bootstrapped predictions, with zero values along the diagonal indicating the absence of self-connections. These visualisations illustrate the robustness of the model estimates and highlight connections exhibiting high uncertainty. **C:** Linear regression fit (Huber robust regression) between original and predicted values of conduction velocity and delay across existing connections.

The bootstrapped delay predictions yielded an average RMSE of 4.38 ± 0.13 ms (95% CI: 4.13 – 4.63 ms), an average MAE of 3.48 ms (95% CI: 3.27 – 3.69 ms) and a median APE of 24%. These results indicate that the model predicts conduction delay with low error and relatively low variability with few connections that exhibit high uncertainty involving regions such as the temporal pole and insula (Figure 7B). For velocity, the robust regression fit showed good correspondence between observed and predicted values, with an accurate fit for mid-range velocities, but slight underestimation at high velocities and slight overestimation at low velocities (Figure 7C). For delay, the model similarly underestimated higher delays and overestimated lower delays (Figure 7C).

## Discussion

### Interaction between white matter microstructure metrics

Our findings revealed coherent relationships among the MRI-derived microstructural parameters. Firstly, axon radius (*r)* showed significant positive correlations with key SMI parameters, including axonal water fraction and fibre alignment (*f* and *p*_2_). In healthy myelinated axons, a higher *f* does not necessarily imply large *r*, elevated *f* values arise either from larger axons, which contain proportionally more intra-axonal water due to their lower surface-to-volume ratio, or from a higher density of small axons (Barakovic et al., 2023; Kunz et al., 2018). Significant positive correlation with *p*_2_ indicates that well-aligned fibres tend to support larger axon diameters (e.g. midbody of the corpus callosum), whereas structures with significant orientation dispersion contain smaller axons (e.g. forceps minor and major; Aboitiz et al., 1992; Zhang et al., 2011), which are potentially more easily tortuous. This is consistent with previous reports that the environment of axons (including the alignment of fibres) influences their morphology, affecting their diameter and overall function (M. Andersson et al., 2020).

Secondly, we found that qT_1_ values were positively correlated with *D*_e,┴_, negatively with *f* and *p*_2_, and showed a weaker negative association with *r*. qT_1_ is inversely related to myelin (strong myelination shortens T_1_) (Callaghan et al., 2015; Stüber et al., 2014). Meanwhile, *D*_e,┴_ will decrease with myelination and increase with demyelination (Guglielmetti et al., 2016; Jelescu et al., 2015, 2016; Liao et al., 2024). As a result, qT_1_ is positively associated with *D*_e,┴_(Travis et al., 2019). Moreover, the negative correlations found between qT_1_ and both *f* and *p*_2_ suggest that voxels with high axonal water fraction and coherent fibre orientation tend to have high myelin volume fraction and thus low qT_1_ (Hutchinson et al., 2024; Jelescu et al., 2016; Klawiter et al., 2011). Since myelin is MR-invisible on diffusion acquisitions due to the short T_2_ of myelin water vs the long TE of the diffusion-weighted spin-echo design (Mackay et al., 1994; Stanisz et al., 1999), the intra-axonal water fraction *f* is a relative weighting of intra- to extra-axonal water, factoring out the myelin volume. Thus, a large myelin volume will indirectly lead to a high *f* by reducing the extra-axonal water volume (Fieremans et al., 2011; Stikov et al., 2015). Our findings indicate that, despite stemming from different MRI contrasts, the parameters effectively capture related microstructural properties of the tissue, particularly concerning axonal organization and myelin content. This cross-validation is particularly important given the distinct sensitivities of the acquisition schemes: multi-shell dMRI targets diffusion properties reflecting cell density and tissue organization (Coelho et al., 2022; Liao et al., 2024; Novikov et al., 2018), the multi-TE diffusion-weighted data emphasizes axonal morphology (Barakovic et al., 2023; Canales-Rodríguez et al., 2024), and qT_1_ primarily reflect myelin density in the WM (Lutti et al., 2014; Piredda, Hilbert, Thiran, et al., 2021; van der Weijden et al., 2021). Together, these complementary measures strengthen the robustness of brain microstructure characterisation and highlight the value of integrating multiple MRI techniques to clarify the relationships between axonal structure, tissue microstructure, myelination, and ultimately conduction velocity in the brain.

### Association of MRI-derived microstructure metrics with CCEP conduction velocity

We combined the F-TRACT fibre delay atlas, which provides cortico-cortical delay data derived from intracerebral depth electrodes (SEEG, Lemaréchal et al., 2022; Trebaul et al., 2018), with fibre length information for the connections between the same pairs of cortical ROIs to yield conduction velocity along these fibres. Since delays were averaged across both propagation directions, the resulting measure is undirected.

We then investigated the association between the aforementioned parameters (*f*, *D*_e,┴_, *p*_2_, *r*, *qT*_1_) and conduction velocity. In line with previous findings (Drakesmith et al., 2019; Ritchie, 1982; Rushton, 1951; Waxman & Bennett, 1972), we observed a strong positive correlation between conduction velocity and *r*, with a significant negative association with qT_1_, confirming that both axon radius and myelin volume are key determinants of conduction velocity in healthy WM. Additionally, among SMI parameters, *f*, *D*_e,┴_, and *p*_2_each showed significant correlations with conduction velocity that were consistent with our hypotheses. For example, a higher *f* is likely associated with larger or more densely packed axons – as discussed previously – that support faster signal propagation. Similarly, lower *D*_e,┴_ suggests a highly tortuous extra-axonal space due to high myelination or high axonal density, also consistent with high conduction velocity (Lobsien et al., 2014). Moreover, a higher *p*_2_ can facilitate more synchronised and efficient action potential transmission along fibre tracts (Clark et al., 2022; Salami et al., 2003). These results support the qualitative agreement of our MRI-derived microstructure parameters with invasive electrophysiological measurements.

Among the studied microstructural parameters, *f* and *r* exhibited the strongest correlation with conduction velocity, followed by *D*_e,┴_, *p*_2_ and qT_1_. Notably, these associations, particularly those involving *f*, qT_1_ and p_2_, were substantially influenced by the length of the fibre (see column *l* in Figure 5). This suggests that fibre length is strongly tied to conduction velocity, as longer fibres typically exhibit larger axon diameters and enhanced structural coherence (myelination) to maintain efficient neural transmission over greater distances (Bajada et al., 2019; Buzsáki et al., 2013; van Blooijs et al., 2023).

To our knowledge, this study is the first to investigate these relationships *in vivo* using diffusion metrics from a clinical MRI system and to validate them against direct conduction velocity measurements.

### White matter conduction velocity and delay prediction: modelling F-TRACT measures from MRI microstructure

While F-TRACT cortico-cortical data provides a direct estimate of axonal conduction delays between subsets of brain regions, these measurements were primarily obtained from patients with epilepsy, during presurgical assessment before resective surgery (Lemaréchal et al., 2022; Trebaul et al., 2018). Therefore, conduction delay (and velocity) cannot be directly measured across the whole brain with this technique and cannot be accessed in healthy volunteers. Previous work used MRI to predict conduction properties in the CC (Berman et al., 2019; Horowitz et al., 2015). More recently, a study by Mancini et al., (2021) proposed a whole-brain estimation of conduction velocity from axon diameter and g-ratio and derived the associated delays using tractography. However, they did not conduct direct validation with ground-truth electrophysiological delay data. In contrast, here we fit a linear model incorporating *r*, *f*, *D*_e,┴_ and qT_1_ to measured delays and velocities in the WM. Our model combining MRI-derived microstructural parameters explains 29% of the variance in measured conduction velocities and enables the extrapolation of conduction velocities along fibre bundles for which no direct electrophysiological measurement is available. None of the four model parameters were redundant. Our model assigned significant positive weights to axon radius, axonal water fraction and qT_1_, and a negative weight to *D*_e,┴_, for predicting conduction velocity.

Given the known biological association between higher myelin content and faster conduction velocities, the positive coefficient on qT_1_ appears counterintuitive. One explanation for this lies in the moderate collinearity observed between qT_1_ and *f*. As mentioned previously, *f* inherently captures variance related not only to axonal packing but also to myelination levels, as these features are often closely coupled (Jelescu et al., 2016). Thus, when both variables are included in the model, axon density captures most of the shared variance related to myelin, leaving qT_1_ to reflect only the remaining, independent variations in tissue properties. Furthermore, although qT_1_ is commonly interpreted as a proxy for myelination, it even captures more variance related to *r*–combination of intra- and extra-axonal compartments (Harkins et al., 2016; Hofer et al., 2015) which could further contribute to its positive association with conduction velocity. As a result, the effect of qT_1_ in the model no longer directly represents myelination alone but a residual signal influenced by other microstructural factors. It is important to note that removing qT1 significantly reduced predicting performances, we thus decided to keep qT1 in the final model. Our findings therefore underscore the importance of carefully interpreting models that include related microstructural measures and demonstrate the value of combining multiple complementary metrics to achieve a more comprehensive understanding of conduction velocity. Future work could employ more specific MRI metrics for myelin content, as those derived from multi-component T_2_ relaxometry (Canales-Rodríguez, Pizzolato, Piredda, et al., 2021; Canales-Rodríguez, Pizzolato, Yu, et al., 2021; Mackay et al., 1994; Piredda, Hilbert, Canales-Rodríguez, et al., 2021).

Our framework linking microstructure to conduction velocity has potential applications in neurological disorders. In conditions such as multiple sclerosis and epilepsy, conduction velocity is often altered by changes in myelination (Sorrentino et al., 2022); since myelination normally ensures rapid, synchronized nerve signalling, myelin abnormalities can disrupt neural communication and promote the hyper-synchronous neuronal firing that underlies seizures (Drenthen et al., 2020; Lassmann, 2014; Poser & Brinar, 2003; Waxman & Bennett, 1972). Similarly, schizophrenia has been linked to subtle myelin abnormalities resulting in disrupted neural synchronisation which can impair whole-brain functional connectivity (Fessel, 2022; Fields, 2008). Comprehensive mapping of conduction delays across the entire brain would thus provide critical insights into brain dynamics and improve models of brain function. Moreover, an independent estimation of conduction velocity of individual WM tracts would augment electroencephalography (EEG) source localisation precision by using priors on delays and could help localise the irritative and the seizure onset zones in patients with epilepsy. Such information could further strengthen recent efforts to predict postsurgical outcome in patients with epilepsy (Mitsuhashi et al., 2021) and could be integrated into computational frameworks such as the Virtual Epileptic Patient model (Makhalova et al., 2022).

Our model predicted conduction velocity with RMSE = 2.60 ± 0.13 ms (median APE = 25%), with only a few connections exhibiting high uncertainty (i.e. inferior-temporal and insula). One key limitation here is the sparse spatial sampling of the SEEG delay data, which restricts the coverage of the whole brain network. In practice, electrodes are placed based on clinical applications rather than uniform coverage. As a result, many areas, notably deep or anatomically secluded cortices like the temporal pole, inferior-temporal and insula, may have few recordings. This inhomogeneous coverage can bias the estimated delay atlas, as connections involving such sparsely sampled regions tend to carry greater uncertainty. Furthermore, even within well-sampled networks, noise and inter-patient variability introduce fluctuations in the delay estimates, meaning that some cortico-cortical connections show greater variability in measured latency than others. This reflects both measurement noise and true biological differences across pathways (Jaber et al., 2024; Lemaréchal et al., 2022). Overall, the restricted SEEG coverage and data heterogeneity impose limits on the atlas’s precision: the method cannot capture delays in unsampled circuits, and delay estimates for sparsely observed connections remain less reliable, prompting caution when interpreting those results.

We need to consider a few limitations for this study. First, dMRI at 7T often yields signal dropouts and spatial distortions in regions with B_0_ field inhomogeneity. These are more pronounced near tissue-air interfaces (e.g. sinuses). In our case, long TE and high *b*-values further impacted the overall data quality in our multi-TE diffusion-weighted acquisition for axon radius estimation, requiring customization of pre-processing tool settings.

Second, the current atlas resolution (Scale 1, 95 ROIs) is relatively coarse. The WM bundle atlas includes four parcellation scales, with increasing numbers of grey matter regions (95, 141, 243, and 473 in scales 1 to 4, respectively). Utilising a finer-grained atlas (e.g., Scale 4, 473 ROIs) could improve our ability to capture variations in microstructure parameters. Additionally, the sample size, particularly for the axon radius estimation, is relatively small. Increasing the number of subjects in the future would enhance the robustness of our findings, especially considering that our delay metric is population-based. Furthermore, the analysis integrates data from different populations: delay and fibre length metrics are derived from population-based atlases (the F-TRACT cohort and HCP tractography), whereas microstructural measures originate from a smaller 7T cohort. Future work can benefit from performing subject-specific tractography in the 7T cohort to directly estimate fibre lengths and improve consistency across measures.

Moreover, combining distinct measurements for conduction delay inherently introduces cumulative uncertainty, as each modality carries its own noise and bias. For example, only for axon diameter estimation the test-retest studies report voxel-wise coefficient of variation of ∼10% for the same metric (Veraart et al., 2021). Likewise, CCEP latency measurements are not perfectly consistent: the N1 peak latency CCEPs typically falls in the 10–50 ms range and can jitter by several milliseconds across trials and subjects (Trebaul et al., 2018). Given all these independent sources of methodological error, our model’s 29% variance explained is a valuable outcome in line with expectations, given the noise floor imposed by the measurements themselves. It is also important to consider limitations and potential improvements related to network properties. Our current predictions are based on an undirected structural connectome derived from dMRI, whereas real physiological signal propagation is directed. Effective connectivity mapping with CCEPs has shown that evoked networks are inherently asymmetric and can therefore reveal directed connections (Crocker et al., 2021; Lemaréchal et al., 2022). Incorporating connection directionality into the model (i.e. distinguishing source and target regions) would likely improve its robustness and biological validity. Moreover, accounting for different path lengths, for example, distinguishing short-range U-fibres from long-range polysynaptic pathways, could further enhance the model.

## Conclusion

In this study, we examined the relationship between conduction velocity, derived from the combination of electrophysiological cortico-cortical delay data with fibre length, and microstructural features obtained from advanced MRI techniques, including axon radius, axonal water fraction, extra-axonal perpendicular diffusivity and quantitative T_1_. Our results highlight significant associations between conduction velocity and these key microstructural parameters. A linear combination of these metrics successfully explained 29% of the variance in conduction velocity, enabling the prediction of whole-brain WM conduction velocities and delays *in vivo*. This combined MRI approach thus offers a promising pathway for characterising neural conduction properties at the whole-brain level non-invasively, with foreseen applications to patients with epilepsy.

## Data and Code Availability

Data and code will be available on a public repository upon article publication.

## Author Contributions

Conceptualization: I.J., P.H.; Data curation: S.A., M.J.; Formal analysis: S.A., Y.A.G., E.J.G. and M.J.; Funding acquisition: I.J., P.H., O.D., S.V. and D.V.D.; Investigation: S.A., H.L., I.R., L.d.W. and E.M.; Methodology: I.J., P.H., S.A., E.J.G., Y.A.G., A.S., T.P., A.C. and M.C. Supervision: I.J., P.H.; Visualization: S.A.; Writing—original draft: S.A.; Writing—review & editing: I.J., A.S., M.J., E.J.G., L.d.W., I.R., H.L., A.C., O.D. and P.H.

## Funding

This study was supported by the Swiss National Science Foundation Sinergia grant no. 209470, Eccellenza Fellowship no. 194260, Swiss Secretariat for Research and Innovation award no. MB22.00032; European Research Council under the European Union’s Seventh Framework Programme (FP/2007–2013)/ERC Grant Agreement no. 616268 F-TRACT, the European Union’s Horizon 2020 Framework Programme for Research and Innovation under Specific Grant Agreement No. 785907 and 945539 (Human Brain Project SGA2 and SGA3), from the Agence Nationale de la Recherche (grant numbers ANR-21-NEUC-0005–01, ANR-22-CE17-0057-03 and ANR-22-PESN-0012).

## Declaration of Competing Interests

None

## Acknowledgments

The authors acknowledge support from the MRI Platform, Fondation Campus Biotech Geneva, Geneva, Switzerland.

## Supplementary Material

### Supplementary Methods

#### SEEG data and axonal conduction delays estimation

The SEEG data analysis was as follows: first, bad channels were identified with a machine learning algorithm supervised by trained personnel. Then, according to standard procedure, the signals were referenced to the bipolar montage to minimize volume conduction effects. Third, short (4 ms) stimulation artefacts were removed by local signal blanking and interpolation. Then, band pass filtering was applied between 1 and 45 Hz and responses to all stimulation pulses within one run were averaged, baseline corrected and z-scored with respect to [−400, −10] ms time before stimulation. Such averaged response was considered significant if its absolute value exceeded a z-score threshold Z=5 within a [0, 80] ms time window. We considered first peak crossing this threshold per signal.

Significant responses were then fitted with a neuronal model according to Dynamic Causal Modelling - an iterative Bayesian procedure of parameters estimation yielding local time constants and axonal delays (Lemaréchal et al., 2022). They were obtained for 651,637 recordings coming from 33,303 stimulations with mean current intensity 3 mA +/− 1 std, mean pulse width 1 ms +/− 0.4 std and frequency 1 Hz. In 73% (27%) of cases stimulations were performed in the biphasic (monophasic) manner. They correspond to 602 implantations (297 males, 301 females, 4 unknown) performed in 567 patients: 276 male (mean age 23 years +/− 14 std), 287 female (mean age 24 years +/− 14 std), 4 unknown (mean age 28 years +/− 10).

## Notes

### Competing Interest Statement

The authors have declared no competing interest.

## References

Aboitiz, F., Scheibel, A. B., Fisher, R. S., & Zaidel, E. (1992a). Fiber composition of the human corpus callosum. Brain Research, 598(1), 143–153. 10.1016/0006-8993(92)90178-C

Aboitiz, F., Scheibel, A. B., Fisher, R. S., & Zaidel, E. (1992b). Fiber composition of the human corpus callosum. Brain Research, 598(1), 143–153. 10.1016/0006-8993(92)90178-C

Alemán-Gómez, Y., Griffa, A., Houde, J.-C., Najdenovska, E., Magon, S., Cuadra, M. B., Descoteaux, M., & Hagmann, P. (2022). A multi-scale probabilistic atlas of the human connectome. Scientific Data, 9(1), 516. 10.1038/s41597-022-01624-8

Alexander, D. C., Dyrby, T. B., Nilsson, M., & Zhang, H. (2019). Imaging brain microstructure with diffusion MRI: Practicality and applications. NMR in Biomedicine, 32(4), e3841. 10.1002/nbm.3841

Alexander, D. C., Hubbard, P. L., Hall, M. G., Moore, E. A., Ptito, M., Parker, G. J. M., & Dyrby, T. B. (2010). Orientationally invariant indices of axon diameter and density from diffusion MRI. NeuroImage, 52(4), 1374–1389. 10.1016/j.neuroimage.2010.05.043

Andersson, J. L. R., Skare, S., & Ashburner, J. (2003). How to correct susceptibility distortions in spin-echo echo-planar images: Application to diffusion tensor imaging. NeuroImage, 20(2), 870–888. 10.1016/S1053-8119(03)00336-7

Andersson, J. L. R., & Sotiropoulos, S. N. (2016). An integrated approach to correction for off-resonance effects and subject movement in diffusion MR imaging. NeuroImage, 125, 1063–1078. 10.1016/j.neuroimage.2015.10.019

Andersson, M., Kjer, H. M., Rafael-Patino, J., Pacureanu, A., Pakkenberg, B., Thiran, J.-P., Ptito, M., Bech, M., Bjorholm Dahl, A., Andersen Dahl, V., & Dyrby, T. B. (2020). Axon morphology is modulated by the local environment and impacts the noninvasive investigation of its structure–function relationship. Proceedings of the National Academy of Sciences, 117(52), 33649–33659. 10.1073/pnas.2012533117

Assaf, Y., Blumenfeld-Katzir, T., Yovel, Y., & Basser, P. J. (2008). Axcaliber: A method for measuring axon diameter distribution from diffusion MRI. Magnetic Resonance in Medicine, 59(6), 1347–1354. 10.1002/mrm.21577

Avalos-Alais, S., Jedynak, M., Boyer, A., Chanteloup-Forêt, B., Pinheiro, C., Cline, C. C., Parmigiani, S., Alemán-Gómez, Y., Hagmann, P., Keller, C. J., David, O., & the F-TRACT Consortium. (2025). High-resolution electrophysiological mapping of effective connectivity of lateral prefrontal cortex. Brain, awaf317. 10.1093/brain/awaf317

Avants, B. B., Tustison, N. J., Song, G., Cook, P. A., Klein, A., & Gee, J. C. (2011). A Reproducible Evaluation of ANTs Similarity Metric Performance in Brain Image Registration. NeuroImage, 54(3), 2033. 10.1016/j.neuroimage.2010.09.025

Bagnato, F., Franco, G., Li, H., Kaden, E., Ye, F., Fan, R., Chen, A., Alexander, D. C., Smith, S. A., Dortch, R., & Xu, J. (2019). Probing axons using multi-compartmental diffusion in multiple sclerosis. Annals of Clinical and Translational Neurology, 6(9), 1595–1605. 10.1002/acn3.50836

Bajada, C. J., Schreiber, J., & Caspers, S. (2019). Fiber length profiling: A novel approach to structural brain organization. NeuroImage, 186, 164–173. 10.1016/j.neuroimage.2018.10.070

Barakovic, M., Pizzolato, M., Tax, C. M. W., Rudrapatna, U., Magon, S., Dyrby, T. B., Granziera, C., Thiran, J.-P., Jones, D. K., & Canales-Rodríguez, E. J. (2023). Estimating axon radius using diffusion-relaxation MRI: Calibrating a surface-based relaxation model with histology. Frontiers in Neuroscience, 17. 10.3389/fnins.2023.1209521

Bartolomei, F., Lagarde, S., Wendling, F., McGonigal, A., Jirsa, V., Guye, M., & Bénar, C. (2017). Defining epileptogenic networks: Contribution of SEEG and signal analysis. Epilepsia, 58(7), 1131–1147. 10.1111/epi.13791

Berman, S., Filo, S., & Mezer, A. A. (2019). Modeling conduction delays in the corpus callosum using MRI-measured g-ratio. NeuroImage, 195, 128–139. 10.1016/j.neuroimage.2019.03.025

Budde, M. D., & Frank, J. A. (2010). Neurite beading is sufficient to decrease the apparent diffusion coefficient after ischemic stroke. Proceedings of the National Academy of Sciences of the United States of America, 107(32), 14472. 10.1073/pnas.1004841107

Buzsáki, G., Logothetis, N., & Singer, W. (2013). Scaling Brain Size, Keeping Timing: Evolutionary Preservation of Brain Rhythms. Neuron, 80(3), 751–764. 10.1016/j.neuron.2013.10.002

Callaghan, M. F., Helms, G., Lutti, A., Mohammadi, S., & Weiskopf, N. (2015). A general linear relaxometry model of R1 using imaging data. Magnetic Resonance in Medicine, 73(3), 1309–1314. 10.1002/mrm.25210

Caminiti, R., Carducci, F., Piervincenzi, C., Battaglia-Mayer, A., Confalone, G., Visco-Comandini, F., Pantano, P., & Innocenti, G. M. (2013). Diameter, Length, Speed, and Conduction Delay of Callosal Axons in Macaque Monkeys and Humans: Comparing Data from Histology and Magnetic Resonance Imaging Diffusion Tractography. Journal of Neuroscience, 33(36), 14501–14511. 10.1523/JNEUROSCI.0761-13.2013

Caminiti, R., Ghaziri, H., Galuske, R., Hof, P. R., & Innocenti, G. M. (2009). Evolution amplified processing with temporally dispersed slow neuronal connectivity in primates. Proceedings of the National Academy of Sciences of the United States of America, 106(46), 19551–19556. 10.1073/pnas.0907655106

Canales-Rodríguez, E. J., Pizzolato, M., Piredda, G. F., Hilbert, T., Kunz, N., Pot, C., Yu, T., Salvador, R., Pomarol-Clotet, E., Kober, T., Thiran, J.-P., & Daducci, A. (2021). Comparison of non-parametric T2 relaxometry methods for myelin water quantification. Medical Image Analysis, 69, 101959. 10.1016/j.media.2021.101959

Canales-Rodríguez, E. J., Pizzolato, M., Yu, T., Piredda, G. F., Hilbert, T., Radua, J., Kober, T., & Thiran, J.-P. (2021). Revisiting the T2 spectrum imaging inverse problem: Bayesian regularized non-negative least squares. NeuroImage, 244, 118582. 10.1016/j.neuroimage.2021.118582

Canales-Rodríguez, E. J., Pizzolato, M., Zhou, F.-L., Barakovic, M., Thiran, J.-P., Jones, D. K., Parker, G. J. M., & Dyrby, T. B. (2024). Pore size estimation in axon-mimicking microfibers with diffusion-relaxation MRI. Magnetic Resonance in Medicine, 91(6), 2579–2596. 10.1002/mrm.29991

Clark, I. A., Mohammadi, S., Callaghan, M. F., & Maguire, E. A. (2022). Conduction velocity along a key white matter tract is associated with autobiographical memory recall ability. eLife, 11, e79303. 10.7554/eLife.79303

Coelho, S., Baete, S. H., Lemberskiy, G., Ades-Aron, B., Barrol, G., Veraart, J., Novikov, D. S., & Fieremans, E. (2022). Reproducibility of the Standard Model of diffusion in white matter on clinical MRI systems. NeuroImage, 257, 119290. 10.1016/j.neuroimage.2022.119290

Coelho, S., Liao, Y., Szczepankiewicz, F., Veraart, J., Chung, S., Lui, Y. W., Novikov, D. S., & Fieremans, E. (2024). Assessment of precision and accuracy of brain white matter microstructure using combined diffusion MRI and relaxometry. Human Brain Mapping, 45(9), e26725. 10.1002/hbm.26725

Crocker, B., Ostrowski, L., Williams, Z. M., Dougherty, D. D., Eskandar, E. N., Widge, A. S., Chu, C. J., Cash, S. S., & Paulk, A. C. (2021). Local and distant responses to single pulse electrical stimulation reflect different forms of connectivity. NeuroImage, 237, 118094. 10.1016/j.neuroimage.2021.118094

David, O., Bastin, J., Chabardès, S., Minotti, L., & Kahane, P. (2010). Studying Network Mechanisms Using Intracranial Stimulation in Epileptic Patients. Frontiers in Systems Neuroscience, 4, 148. 10.3389/fnsys.2010.00148

Dong, J. W., Jelescu, I. O., Ades-Aron, B., Novikov, D. S., Friedman, K., Babb, J. S., Osorio, R. S., Galvin, J. E., Shepherd, T. M., & Fieremans, E. (2020). Diffusion MRI biomarkers of white matter microstructure vary nonmonotonically with increasing cerebral amyloid deposition. Neurobiology of Aging, 89, 118–128. 10.1016/j.neurobiolaging.2020.01.009

Drakesmith, M., Harms, R., Rudrapatna, S. U., Parker, G. D., Evans, C. J., & Jones, D. K. (2019). Estimating axon conduction velocity in vivo from microstructural MRI. NeuroImage, 203, 116186. 10.1016/j.neuroimage.2019.116186

Drenthen, G. S., Backes, W. H., Aldenkamp, A. P., Vermeulen, R. J., Klinkenberg, S., & Jansen, J. F. A. (2020). On the merits of non-invasive myelin imaging in epilepsy, a literature review. Journal of Neuroscience Methods, 338, 108687. 10.1016/j.jneumeth.2020.108687

Duval, T., McNab, J. A., Setsompop, K., Witzel, T., Schneider, T., Huang, S. Y., Keil, B., Klawiter, E. C., Wald, L. L., & Cohen-Adad, J. (2015). In vivo mapping of human spinal cord microstructure at 300 mT/m. Neuroimage, 118, 494–507.

Dyrby, T. B., Søgaard, L. V., Hall, M. G., Ptito, M., & Alexander, D. C. (2013). Contrast and stability of the axon diameter index from microstructure imaging with diffusion MRI. Magnetic Resonance in Medicine, 70(3), 711–721. 10.1002/mrm.24501

Fan, Q., Tian, Q., Ohringer, N. A., Nummenmaa, A., Witzel, T., Tobyne, S. M., Klawiter, E. C., Mekkaoui, C., Rosen, B. R., Wald, L. L., Salat, D. H., & Huang, S. Y. (2019). Age-related alterations in axonal microstructure in the corpus callosum measured by high-gradient diffusion MRI. NeuroImage, 191, 325. 10.1016/j.neuroimage.2019.02.036

Feher, J. (2012). 3.3—Propagation of the Action Potential. In J. Feher (Ed.), Quantitative Human Physiology (pp. 227–235). Academic Press. 10.1016/B978-0-12-382163-8.00025-6

Fessel, J. (2022). Abnormal oligodendrocyte function in schizophrenia explains the long latent interval in some patients. Translational Psychiatry, 12(1), 120. 10.1038/s41398-022-01879-0

Fields, R. D. (2008). White matter in learning, cognition and psychiatric disorders. Trends in Neurosciences, 31(7), 361–370. 10.1016/j.tins.2008.04.001

Fieremans, E., Jensen, J. H., & Helpern, J. A. (2011). White matter characterization with diffusional kurtosis imaging. NeuroImage, 58(1), 177–188. 10.1016/j.neuroimage.2011.06.006

Guglielmetti, C., Veraart, J., Roelant, E., Mai, Z., Daans, J., Van Audekerke, J., Naeyaert, M., Vanhoutte, G., Delgado y Palacios, R., Praet, J., Fieremans, E., Ponsaerts, P., Sijbers, J., Van der Linden, A., & Verhoye, M. (2016). Diffusion kurtosis imaging probes cortical alterations and white matter pathology following cuprizone induced demyelination and spontaneous remyelination. NeuroImage, 125, 363–377. 10.1016/j.neuroimage.2015.10.052

Hagmann, P., Cammoun, L., Gigandet, X., Meuli, R., Honey, C. J., Wedeen, V. J., & Sporns, O. (2008). Mapping the Structural Core of Human Cerebral Cortex. PLOS Biology, 6(7), e159. 10.1371/journal.pbio.0060159

Harkins, K. D., Xu, J., Dula, A. N., Li, K., Valentine, W. M., Gochberg, D. F., Gore, J. C., & Does, M. D. (2016). The Microstructural Correlates of T1 in White Matter. Magnetic Resonance in Medicine, 75(3), 1341–1345. 10.1002/mrm.25709

Hartline, D. K., & Colman, D. R. (2007). Rapid conduction and the evolution of giant axons and myelinated fibers. Current Biology: CB, 17(1), R29–35. 10.1016/j.cub.2006.11.042

Hofer, S., Wang, X., Roeloffs, V., & Frahm, J. (2015). Single-shot T1 mapping of the corpus callosum: A rapid characterization of fiber bundle anatomy. Frontiers in Neuroanatomy, 9, 57. 10.3389/fnana.2015.00057

Horowitz, A., Barazany, D., Tavor, I., Bernstein, M., Yovel, G., & Assaf, Y. (2015). In vivo correlation between axon diameter and conduction velocity in the human brain. Brain Structure and Function, 220(3), 1777–1788. 10.1007/s00429-014-0871-0

Hursh, J. B. (1939). CONDUCTION VELOCITY AND DIAMETER OF NERVE FIBERS. American Journal of Physiology-Legacy Content, 127(1), 131–139. 10.1152/ajplegacy.1939.127.1.131

Hutchinson, G., Thotland, J., Pisharady, P. K., Garwood, M., Lenglet, C., & Kauppinen, R. A. (2024). T1 relaxation and axon fibre configuration in human white matter. NMR in Biomedicine, 37(12), e5234. 10.1002/nbm.5234

Innocenti, G. M., Vercelli, A., & Caminiti, R. (2014). The Diameter of Cortical Axons Depends Both on the Area of Origin and Target. Cerebral Cortex, 24(8), 2178–2188. 10.1093/cercor/bht070

Jaber, K., Avigdor, T., Mansilla, D., Ho, A., Thomas, J., Abdallah, C., Chabardes, S., Hall, J., Minotti, L., Kahane, P., Grova, C., Gotman, J., & Frauscher, B. (2024). A spatial perturbation framework to validate implantation of the epileptogenic zone. Nature Communications, 15, 5253. 10.1038/s41467-024-49470-z

Jedynak, M., Boyer, A., Lemaréchal, J.-D., Trebaul, L., Tadel, F., Bhattacharjee, M., Chanteloup-Forêt, B., Deman, P., Tuyisenge, V., Ayoubian, L., Hugues, E., Saubat-Guigui, C., Zouglech, R., Reyes-Mejia, G. C., Tourbier, S., Hagmann, P., Adam, C., Barba, C., Bartolomei, F., … F-TRACT Consortium. (2023). F-TRACT: a probabilistic atlas of anatomo-functional connectivity of the human brain (F-TRACT_P_11_v2307). EBRAINS. 10.25493/5AM4-J3F

Jelescu, I. O., & Fieremans, E. (2023). Chapter 2—Sensitivity and specificity of diffusion MRI to neuroinflammatory processes. In C. Laule & J. D. Port (Eds.), Advances in Magnetic Resonance Technology and Applications (Vol. 9, pp. 31–50). Academic Press. 10.1016/B978-0-323-91771-1.00010-1

Jelescu, I. O., Palombo, M., Bagnato, F., & Schilling, K. G. (2020). Challenges for biophysical modeling of microstructure. Journal of Neuroscience Methods, 344, 108861. 10.1016/j.jneumeth.2020.108861

Jelescu, I. O., Veraart, J., Adisetiyo, V., Milla, S. S., Novikov, D. S., & Fieremans, E. (2015). One diffusion acquisition and different white matter models: How does microstructure change in human early development based on WMTI and NODDI? NeuroImage, 107, 242–256. 10.1016/j.neuroimage.2014.12.009

Jelescu, I. O., Zurek, M., Winters, K. V., Veraart, J., Rajaratnam, A., Kim, N. S., Babb, J. S., Shepherd, T. M., Novikov, D. S., Kim, S. G., & Fieremans, E. (2016). *In vivo* quantification of demyelination and recovery using compartment-specific diffusion MRI metrics validated by electron microscopy. NeuroImage, 132, 104–114. 10.1016/j.neuroimage.2016.02.004

Kellner, E., Dhital, B., Kiselev, V. G., & Reisert, M. (2016). Gibbs-ringing artifact removal based on local subvoxel-shifts. Magnetic Resonance in Medicine, 76(5), 1574–1581. 10.1002/mrm.26054

Klawiter, E. C., Schmidt, R. E., Trinkaus, K., Liang, H.-F., Budde, M. D., Naismith, R. T., Song, S.-K., Cross, A. H., & Benzinger, T. L. (2011). Radial Diffusivity Predicts Demyelination in ex-vivo Multiple Sclerosis Spinal Cords. NeuroImage, 55(4), 1454–1460. 10.1016/j.neuroimage.2011.01.007

Koenig, S. H., Brown III, R. D., Spiller, M., & Lundbom, N. (1990). Relaxometry of brain: Why white matter appears bright in MRI. Magnetic Resonance in Medicine, 14(3), 482–495. 10.1002/mrm.1910140306

Kunz, N., da Silva, A. R., & Jelescu, I. O. (2018). Intra- and extra-axonal axial diffusivities in the white matter: Which one is faster? NeuroImage, 181, 314–322. 10.1016/j.neuroimage.2018.07.020

Lassmann, H. (2014). Mechanisms of white matter damage in multiple sclerosis. Glia, 62(11), 1816–1830. 10.1002/glia.22597

Lee, H.-H., Papaioannou, A., Kim, S.-L., Novikov, D. S., & Fieremans, E. (2020). A time-dependent diffusion MRI signature of axon caliber variations and beading. Communications Biology, 3(1), 1–13. 10.1038/s42003-020-1050-x

Lemaréchal, J.-D., Jedynak, M., Trebaul, L., Boyer, A., Tadel, F., Bhattacharjee, M., Deman, P., Tuyisenge, V., Ayoubian, L., Hugues, E., Chanteloup-Forêt, B., Saubat, C., Zouglech, R., Mejia, G. C. R., Tourbier, S., Hagmann, P., Adam, C., Barba, C., Bartolomei, F., … Consortium, F.-T. (2022). A brain atlas of axonal and synaptic delays based on modelling of cortico-cortical evoked potentials. Brain, 145(5), 1653. 10.1093/brain/awab362

Liao, Y., Coelho, S., Chen, J., Ades-Aron, B., Pang, M., Stepanov, V., Osorio, R., Shepherd, T., Lui, Y. W., Novikov, D. S., & Fieremans, E. (2024). Mapping tissue microstructure of brain white matter in vivo in health and disease using diffusion MRI. Imaging Neuroscience, 2, 1–17. 10.1162/imag_a_00102

Liewald, D., Miller, R., Logothetis, N., Wagner, H.-J., & Schüz, A. (2014). Distribution of axon diameters in cortical white matter: An electron-microscopic study on three human brains and a macaque. Biological Cybernetics, 108(5), 541–557. 10.1007/s00422-014-0626-2

Lobsien, D., Ettrich, B., Sotiriou, K., Classen, J., Then Bergh, F., & Hoffmann, K.-T. (2014). Whole-Brain Diffusion Tensor Imaging in Correlation to Visual-Evoked Potentials in Multiple Sclerosis: A Tract-Based Spatial Statistics Analysis. AJNR: American Journal of Neuroradiology, 35(11), 2076–2081. 10.3174/ajnr.A4034

Lutti, A., Dick, F., Sereno, M. I., & Weiskopf, N. (2014). Using high-resolution quantitative mapping of R1 as an index of cortical myelination. NeuroImage, 93, 176–188. 10.1016/j.neuroimage.2013.06.005

Mackay, A., Whittall, K., Adler, J., Li, D., Paty, D., & Graeb, D. (1994). In vivo visualization of myelin water in brain by magnetic resonance. Magnetic Resonance in Medicine, 31(6), 673–677. 10.1002/mrm.1910310614

Makhalova, J., Medina Villalon, S., Wang, H., Giusiano, B., Woodman, M., Bénar, C., Guye, M., Jirsa, V., & Bartolomei, F. (2022). Virtual epileptic patient brain modeling: Relationships with seizure onset and surgical outcome. Epilepsia, 63(8), 1942–1955. 10.1111/epi.17310

Mancini, M., Tian, Q., Fan, Q., Cercignani, M., & Huang, S. Y. (2021). Dissecting whole-brain conduction delays through MRI microstructural measures. Brain Structure and Function, 226(8), 2651–2663. 10.1007/s00429-021-02358-w

Mezer, A., Yeatman, J. D., Stikov, N., Kay, K. N., Cho, N.-J., Dougherty, R. F., Perry, M. L., Parvizi, J., Hua, L. H., Butts-Pauly, K., & Wandell, B. A. (2013). Quantifying the local tissue volume and composition in individual brains with magnetic resonance imaging. Nature Medicine, 19(12), 1667–1672. 10.1038/nm.3390

Mitsuhashi, T., Sonoda, M., Sakakura, K., Jeong, J.-W., Luat, A. F., Sood, S., & Asano, E. (2021). Dynamic tractography-based localization of spike sources and animation of spike propagations. Epilepsia, 62(10), 2372–2384. 10.1111/epi.17025

Moeller, S., Pisharady, P. K., Ramanna, S., Lenglet, C., Wu, X., Dowdle, L., Yacoub, E., Uğurbil, K., & Akçakaya, M. (2021). NOise reduction with DIstribution Corrected (NORDIC) PCA in dMRI with complex-valued parameter-free locally low-rank processing. NeuroImage, 226, 117539. 10.1016/j.neuroimage.2020.117539

Moore, J. W., Joyner, R. W., Brill, M. H., Waxman, S. D., & Najar-Joa, M. (1978). Simulations of conduction in uniform myelinated fibers. Relative sensitivity to changes in nodal and internodal parameters. Biophysical Journal, 21(2), 147–160.

Novikov, D. S., Fieremans, E., Jespersen, S. N., & Kiselev, V. G. (2019). Quantifying brain microstructure with diffusion MRI: Theory and parameter estimation. NMR in Biomedicine, 32(4), e3998. 10.1002/nbm.3998

Novikov, D. S., Veraart, J., Jelescu, I. O., & Fieremans, E. (2018). Rotationally-invariant mapping of scalar and orientational metrics of neuronal microstructure with diffusion MRI. NeuroImage, 174, 518–538. 10.1016/j.neuroimage.2018.03.006

Pavan, T., Alemán-Gómez, Y., Jenni, R., Steullet, P., Schilliger, Z., Dwir, D., Cleusix, M., Alameda, L., Do, K. Q., Conus, P., Klauser, P., Hagmann, P., & Jelescu, I. (2025). White Matter Microstructure Alterations in Early Psychosis and Schizophrenia (p. 2024.02.01.24301979). medRxiv. 10.1101/2024.02.01.24301979

Piredda, G. F., Hilbert, T., Canales-Rodríguez, E. J., Pizzolato, M., von Deuster, C., Meuli, R., Pfeuffer, J., Daducci, A., Thiran, J.-P., & Kober, T. (2021). Fast and high-resolution myelin water imaging: Accelerating multi-echo GRASE with CAIPIRINHA. Magnetic Resonance in Medicine, 85(1), 209–222. 10.1002/mrm.28427

Piredda, G. F., Hilbert, T., Thiran, J.-P., & Kober, T. (2021). Probing myelin content of the human brain with MRI: A review. Magnetic Resonance in Medicine, 85(2), 627–652. 10.1002/mrm.28509

Poser, C. M., & Brinar, V. V. (2003). Epilepsy and multiple sclerosis. Epilepsy & Behavior, 4(1), 6–12. 10.1016/S1525-5050(02)00646-7

Purves, D., Augustine, G. J., Fitzpatrick, D., Katz, L. C., LaMantia, A.-S., McNamara, J. O., & Williams, S. M. (2001). Increased Conduction Velocity as a Result of Myelination. In Neuroscience. 2nd edition. Sinauer Associates. https://www.ncbi.nlm.nih.gov/books/NBK10921/

Ritchie, J. M. (1982). On the relation between fibre diameter and conduction velocity in myelinated nerve fibres. Proceedings of the Royal Society of London. Series B, Biological Sciences, 217(1206), 29–35. 10.1098/rspb.1982.0092

Rushton, W. A. H. (1951). A theory of the effects of fibre size in medullated nerve. The Journal of Physiology, 115(1), 101–122. 10.1113/jphysiol.1951.sp004655

Salami, M., Itami, C., Tsumoto, T., & Kimura, F. (2003). Change of conduction velocity by regional myelination yields constant latency irrespective of distance between thalamus and cortex. Proceedings of the National Academy of Sciences, 100(10), 6174–6179. 10.1073/pnas.0937380100

Sorrentino, P., Petkoski, S., Sparaco, M., Lopez, E. T., Signoriello, E., Baselice, F., Bonavita, S., Pirozzi, M. A., Quarantelli, M., Sorrentino, G., & Jirsa, V. (2022). Whole-Brain Propagation Delays in Multiple Sclerosis, a Combined Tractography-Magnetoencephalography Study. Journal of Neuroscience, 42(47), 8807–8816. 10.1523/JNEUROSCI.0938-22.2022

Stanisz, G. j., Kecojevic, A., Bronskill, M. j., & Henkelman, R. m. (1999). Characterizing white matter with magnetization transfer and T2. Magnetic Resonance in Medicine, 42(6), 1128–1136. 10.1002/(SICI)1522-2594(199912)42:6%253C1128::AID-MRM18%253E3.0.CO;2-9

Stikov, N., Campbell, J. S. W., Stroh, T., Lavelée, M., Frey, S., Novek, J., Nuara, S., Ho, M.-K., Bedell, B. J., Dougherty, R. F., Leppert, I. R., Boudreau, M., Narayanan, S., Duval, T., Cohen-Adad, J., Picard, P.-A., Gasecka, A., Côté, D., & Pike, G. B. (2015). In vivo histology of the myelin g-ratio with magnetic resonance imaging. NeuroImage, 118, 397–405. 10.1016/j.neuroimage.2015.05.023

Stüber, C., Morawski, M., Schäfer, A., Labadie, C., Wähnert, M., Leuze, C., Streicher, M., Barapatre, N., Reimann, K., Geyer, S., Spemann, D., & Turner, R. (2014). Myelin and iron concentration in the human brain: A quantitative study of MRI contrast. NeuroImage, 93, 95–106. 10.1016/j.neuroimage.2014.02.026

Swadlow, H. A., Rosene, D. L., & Waxman, S. G. (1978). Characteristics of interhemispheric impulse conduction between prelunate gyri of the rhesus monkey. Experimental Brain Research, 33(3), 455–467. 10.1007/BF00235567

Tomlinson, S. B., Baumgartner, M. E., Darlington, T. R., Marsh, E. D., & Kennedy, B. C. (2025). Mapping the Epileptogenic Brain Using Low-Frequency Stimulation: Two Decades of Advances and Uncertainties. Journal of Clinical Medicine, 14(6), 1956. 10.3390/jcm14061956

Toschi, N., Gisbert, R. A., Passamonti, L., Canals, S., & De Santis, S. (2020). Multishell diffusion imaging reveals sex-specific trajectories of early white matter degeneration in normal aging. Neurobiology of Aging, 86, 191–200. 10.1016/j.neurobiolaging.2019.11.014

Tournier, J.-D., Smith, R., Raffelt, D., Tabbara, R., Dhollander, T., Pietsch, M., Christiaens, D., Jeurissen, B., Yeh, C.-H., & Connelly, A. (2019). MRtrix3: A fast, flexible and open software framework for medical image processing and visualisation. NeuroImage, 202, 116137. 10.1016/j.neuroimage.2019.116137

Travis, K. E., Castro, M. R. H., Berman, S., Dodson, C. K., Mezer, A. A., Ben-Shachar, M., & Feldman, H. M. (2019). More than myelin: Probing white matter differences in prematurity with quantitative T1 and diffusion MRI. NeuroImage : Clinical, 22, 101756. 10.1016/j.nicl.2019.101756

Trebaul, L., Deman, P., Tuyisenge, V., Jedynak, M., Hugues, E., Rudrauf, D., Bhattacharjee, M., Tadel, F., Chanteloup-Foret, B., Saubat, C., Reyes Mejia, G. C., Adam, C., Nica, A., Pail, M., Dubeau, F., Rheims, S., Trébuchon, A., Wang, H., Liu, S., … David, O. (2018). Probabilistic functional tractography of the human cortex revisited. NeuroImage, 181, 414–429. 10.1016/j.neuroimage.2018.07.039

van Blooijs, D., van den Boom, M. A., van der Aar, J. F., Huiskamp, G. M., Castegnaro, G., Demuru, M., Zweiphenning, W. J. E. M., van Eijsden, P., Miller, K. J., Leijten, F. S. S., & Hermes, D. (2023). Developmental trajectory of transmission speed in the human brain. Nature Neuroscience, 26(4), 537–541. 10.1038/s41593-023-01272-0

van der Weijden, C. W. J., García, D. V., Borra, R. J. H., Thurner, P., Meilof, J. F., van Laar, P.-J., Dierckx, R. A. J. O., Gutmann, I. W., & de Vries, E. F. J. (2021). Myelin quantification with MRI: A systematic review of accuracy and reproducibility. NeuroImage, 226, 117561. 10.1016/j.neuroimage.2020.117561

Van Essen, D. C., Smith, S. M., Barch, D. M., Behrens, T. E. J., Yacoub, E., & Ugurbil, K. (2013). The WU-Minn Human Connectome Project: An overview. NeuroImage, 80, 62–79. 10.1016/j.neuroimage.2013.05.041

Veraart, J., Fieremans, E., & Novikov, D. S. (2019). On the scaling behavior of water diffusion in human brain white matter. NeuroImage, 185, 379–387. 10.1016/j.neuroimage.2018.09.075

Veraart, J., Novikov, D. S., & Fieremans, E. (2018). TE dependent Diffusion Imaging (TEdDI) distinguishes between compartmental T2 relaxation times. NeuroImage, 182, 360–369. 10.1016/j.neuroimage.2017.09.030

Veraart, J., Nunes, D., Rudrapatna, U., Fieremans, E., Jones, D. K., Novikov, D. S., & Shemesh, N. (2020). Noninvasive quantification of axon radii using diffusion MRI. eLife, 9, e49855. 10.7554/eLife.49855

Veraart, J., Raven, E. P., Edwards, L. J., Weiskopf, N., & Jones, D. K. (2021). The variability of MR axon radii estimates in the human white matter. Human Brain Mapping, 42(7), 2201–2213. 10.1002/hbm.25359

Vogt, N. M., Hunt, J. F., Adluru, N., Dean, D. C., III, Johnson, S. C., Asthana, S., Yu, J.-P. J., Alexander, A. L., & Bendlin, B. B. (2020). Cortical Microstructural Alterations in Mild Cognitive Impairment and Alzheimer’s Disease Dementia. Cerebral Cortex, 30(5), 2948–2960. 10.1093/cercor/bhz286

Waxman, S. G. (1980). Determinants of conduction velocity in myelinated nerve fibers. Muscle & Nerve, 3(2), 141–150. 10.1002/mus.880030207

Waxman, S. G., & Bennett, M. V. L. (1972). Relative Conduction Velocities of Small Myelinated and Non-myelinated Fibres in the Central Nervous System. Nature New Biology, 238(85), 217–219. 10.1038/newbio238217a0

Wendling, F., Chauvel, P., Biraben, A., & Bartolomei, F. (2010). From Intracerebral EEG Signals to Brain Connectivity: Identification of Epileptogenic Networks in Partial Epilepsy. Frontiers in Systems Neuroscience, 4. 10.3389/fnsys.2010.00154

Xie, M., Tobin, J. E., Budde, M. D., Chen, C.-I., Trinkaus, K., Cross, A. H., McDaniel, D. P., Song, S.-K., & Armstrong, R. C. (2010). Rostrocaudal Analysis of Corpus Callosum Demyelination and Axon Damage Across Disease Stages Refines Diffusion Tensor Imaging Correlations With Pathological Features. Journal of Neuropathology & Experimental Neurology, 69(7), 704–716. 10.1097/NEN.0b013e3181e3de90

Zhang, H., Hubbard, P. L., Parker, G. J. M., & Alexander, D. C. (2011). Axon diameter mapping in the presence of orientation dispersion with diffusion MRI. NeuroImage, 56(3), 1301–1315. 10.1016/j.neuroimage.2011.01.084

